# Benchmarking Large Language Models for Bio-Image Analysis Code Generation

**DOI:** 10.1101/2024.04.19.590278

**Authors:** Robert Haase, Christian Tischer, Jean-Karim Hériché, Nico Scherf

**Affiliations:** Data Science Center, Leipzig University, Humboldtstraße 25, 04105 Leipzig, Germany; Center for Scalable Data Analytics and Artificial Intelligence (ScaDS.AI) Dresden / Leipzig; Data Science Centre, European Molecular Biology Laboratory, Meyerhofstraße 1, 69117 Heidelberg, Germany; Cell Biology and Biophysics Unit, European Molecular Biology Laboratory, Meyerhofstraße 1, 69117 Heidelberg, Germany; Max Planck Institute for Human Cognitive and Brain Sciences, Stephanstraße 1A, 04103, Leipzig, Germany

## Abstract

In the computational age, life-scientists often have to write Python code to solve bio-image analysis (BIA) problems. Many of them have not been formally trained in programming though. Code-generation, or coding assistance in general, with Large Language Models (LLMs) can have a clear impact on BIA. To the best of our knowledge, the quality of the generated code in this domain has not been studied. We present a quantitative benchmark to estimate the capability of LLMs to generate code for solving common BIA tasks. Our benchmark currently consists of 57 human-written prompts with corresponding reference solutions in Python, and unit-tests to evaluate functional correctness of potential solutions. We demonstrate our benchmark here and compare 18 state-of-the-art LLMs. To ensure that we will cover most of our community needs we also outline mid- and long-term strategies to maintain and extend the benchmark by the BIA open-source community. This work should support users in deciding for an LLM and also guide LLM developers in improving the capabilities of LLMs in the BIA domain.

## 1 Introduction

Many projects in biology involve state-of-the-art microscopy and quantitative bio-image analysis (BIA), which increasingly requires solid programming skills for the experimentalists. As programming is commonly not taught to life-scientists, we see potential in using large language models to assist people in this task. Modern Large Language Models (LLMs) such as chatGPT (OpenAI et al. 2023) change the way how humans interact with computers. LLMs were originally developed to solve natural language processing tasks such as text classification, language translation, or question answering. These models are also capable of translating human languages into programming languages, e.g. from English to Python. They can produce executable code that solves a task defined by human natural language input [6]. This capability has huge potential for interdisciplinary research areas such as microscopy bio-image analysis [22]. LLMs can fill a gap where scientists with limited programming skills meet more advanced image analysis tasks. LLMs are indeed capable of writing BIA code as demonstrated in [23], but it is yet unclear where the limitations of this technology are in the BIA context. So a systematic way to analyze LLMs in this domain is needed. In a more general setting multiple LLM code generation benchmarks have been proposed [7,3,17,27,13]. We think the bioimaging community urgently needs its own benchmark, an openly accessible, quantitative way to measure LLM capabilities, in particular given that LLM technology is developing rapidly. Here, we present the core of this benchmark. It is derived from HumanEval [7], an established code-generation benchmark and tailored for scientific data analysis questions in the bioimaging context.

All code used for the benchmark, sampled prompt-responses from the evaluated LLMs, and Python Jupyter notebooks for reproducing Figures in this preprint are available via this Github repository: https://github.com/haesleinhuepf/human-eval-bia

## 2 Methods

Our benchmark currently consists of 57 human-written Python functions containing documentation, the docstring, of what the function is supposed to do. An example is shown in Figure 1. We kept the docstring intentionally brief and natural, because we intend to use LLMs to facilitate coding for bio-image analysts and this would better reflect a typical use-case. The docstring and the function signature is then passed to an LLM as part of a prompt asking to complete the code. Note that the human-written implementation function body just serves as a reference solution and is not passed to the LLM. Our benchmark also provides the unit-tests for the functions, so we can evaluate functional correctness. If the code generated by the language model is executable and produces results which pass these unit-tests, we consider the LLM has having solved the problem. Every prompt is sent multiple times to the LLM, and we track how often the generated code passes the tests. Here, we follow the established standard measure pass@k [7] estimating the probability that, if asked k times, the LLM will at least once give a correct answer. We particularly focus on the practically most relevant special case pass@1, i.e. we want to know how likely it is that the first generated solution works. We also summarize the libraries required by the generated code and the typical resulting error messages.

**Figure 1.**
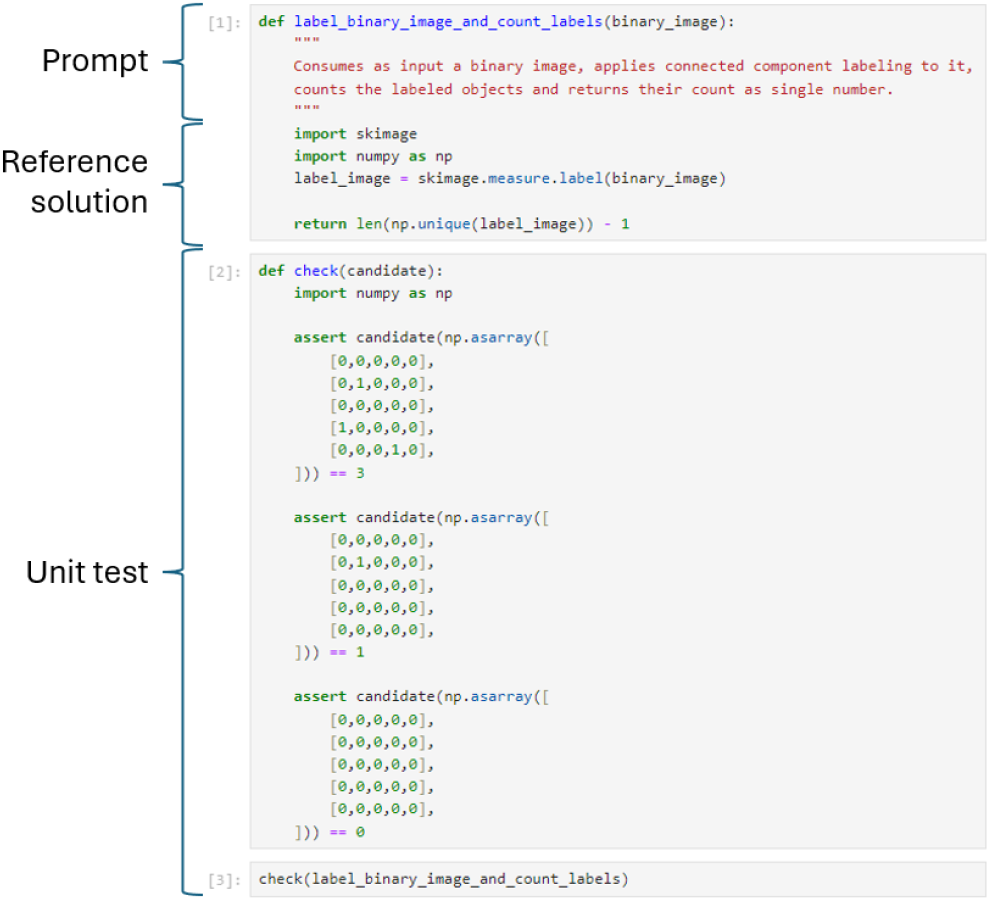
The example test-case label_binary_image_and_count_labels is implemented as Jupyter Notebook consisting of a function signature consuming a binary image and a docstring describing what the function is supposed to do. These two serve as part of a prompt to the LLM asking to complete the code. The function body serves as a reference solution (sometimes referred to as canonical solution), which allows writing a unit test. If the unit test passes for the code generated by an LLM we define the LLM as capable of solving the task. An up-to-date list of all test-cases is available online:https://github.com/haesleinhuepf/human-eval-bia/blob/main/test_cases/readme.md

Our selected prompts range from basic image analysis tasks, such as applying an edge-preserving denoising filter to an image, over intermediate tasks such as labeling objects in a binary image and counting them, shown in Figure 1, to more challenging workflows combining image processing steps, descriptive statistics, tabular data wrangling and dimensionality reduction. There is also a positive-control test-case, called return_hello_world, which is intentionally kept very simple to test if a specific LLM model is capable of solving a trivial base task at all.

To enable extension of our benchmark and reproduction of our results, we provide the infrastructure to turn the folder of test-case Jupyter Notebooks into a JSONL file suitable for evaluation with HumanEval [7]. We also did minor modifications to this framework to be able to execute the benchmark for our purposes. For example, we added code that moves example images to the temporary folder from which the test-case code is executed. With this, our benchmark can cover functions that require accessing files and folders, which the original HumanEval benchmark was not capable of. All modifications are explained in our Github repository and the supplementary Zip file.

We introduce our benchmark by comparing the capabilities of a range of state-of-the-art LLMs covering commercial and freely available or open source models. We cover gemini-pro [25], gemini-1.5-flash-001, gpt-3.5-turbo-1106, gpt-4-1106-preview, gpt-4-2024-04-09, gpt-4o-2024-05-13, codegemma, codellama [24], claude-3-opus-20240229 [2], claude-3-5-sonnet-20240620, command-r-plus [8], llama3 [21], mixtral [16] and phi3 [1]. The gemini-pro model was accessed via the Google Vertex API [14], which did not support specifying a model version. Thus, we document here that the benchmark was executed on April 16th and 17th 2024. Code for benchmarking gemini-1.5-pro and gemini-ultra are available as well, but we were not able to execute it due to rate limits. For the open source models, we set up two kubernetes clusters each with 128 GB of RAM and 4 GPUs (one cluster with Tesla P40 and one with RTX 2080) running ollama version 0.1.32 [19]. The open source models versions were codegemma:7b-instruct-fp16, codellama:70b-instruct-q4, command-r-plus:104b-q4, llama3:70b-instruct-q8, llama3:70b-instruct-q4, llama3:8b-instruct-fp16, mixtral:8×22b-instruct-v0.1-q4, mixtral:8×7b-instruct-v0.1-q5 and phi3:3.8b-mini-instruct-4k-fp16. The open source model codellama:7b-instruct-q4, named codellama in the following and in Figures for technical reasons, was prompted using ollama version 0.1.29 for Windows [20].

To benchmark the models, we generated 10 code samples for each of the 57 test-cases from each of the 18 models. Benchmarking of the commercial models was done on a Windows 10 Laptop with an AMD Ryzen 9 6900 CPU, 32 GB of RAM and a NVidia RTX 3050 TI GPU with 4 GB of RAM. For the open source models, the notebooks were run on a virtual machine with Intel Xeon Gold 6226R CPUs, 24 GB of RAM and a NVIDIA V100S-8Q GPU with 8GB of RAM.

All test-cases (human-readable Jupyter Notebooks and packaged as JSONL file), sampling and evaluation code, generated samples and data analysis/visualization notebooks are available in the Github repository of the project. All respective Python package versions are documented in the environment.yml file in the Github repository and the supplementary Zip file facilitating full reproducibility of our analysis.

## 3 Results

The pass-rates visualized in Figure 2 correspond to pass@1 counting the success rate from drawn examples. Detailed pass@k rates with k=1, k=5 and k=10 are shown in Figure 3. They reveal that the three leading models, claude-3-5-sonnet-20240620, gpt-4o-2024-05-13 and gpt-4-turbo-2024-04-09 have a similar performance in terms of pass-rates of 58 ± 40%, 50 ± 41% and 47 ± 38%, respectively.Analysis of the pass-rates for individual test-cases are shown in Figure 4. The results highlight that most of the test-cases were solved by at least one LLM. Interestingly, some test-cases could not be solved by any LLM, even though we would consider them relatively simple, e.g. deconvolve_image, extract_surface_measure_area and open_image_read_voxel_size.

**Figure 2.**
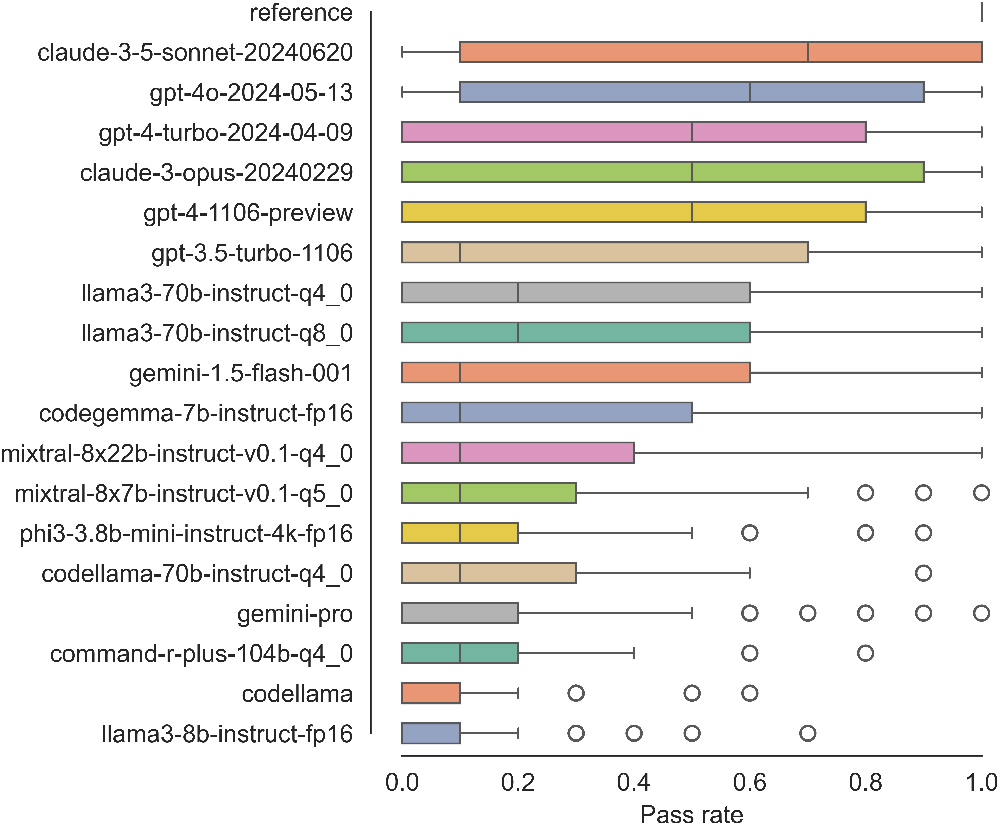
Quantitative pass-rate comparison of all tested LLMs and, as a sanity check, the human reference solution: Measured fraction of passed tests visualized as box plot summarizing measurements from 57 test-cases. The corresponding, updated notebook is available online:https://github.com/haesleinhuepf/human-eval-bia/blob/main/demo/summarize_by_case.ipynb

**Figure 3.**
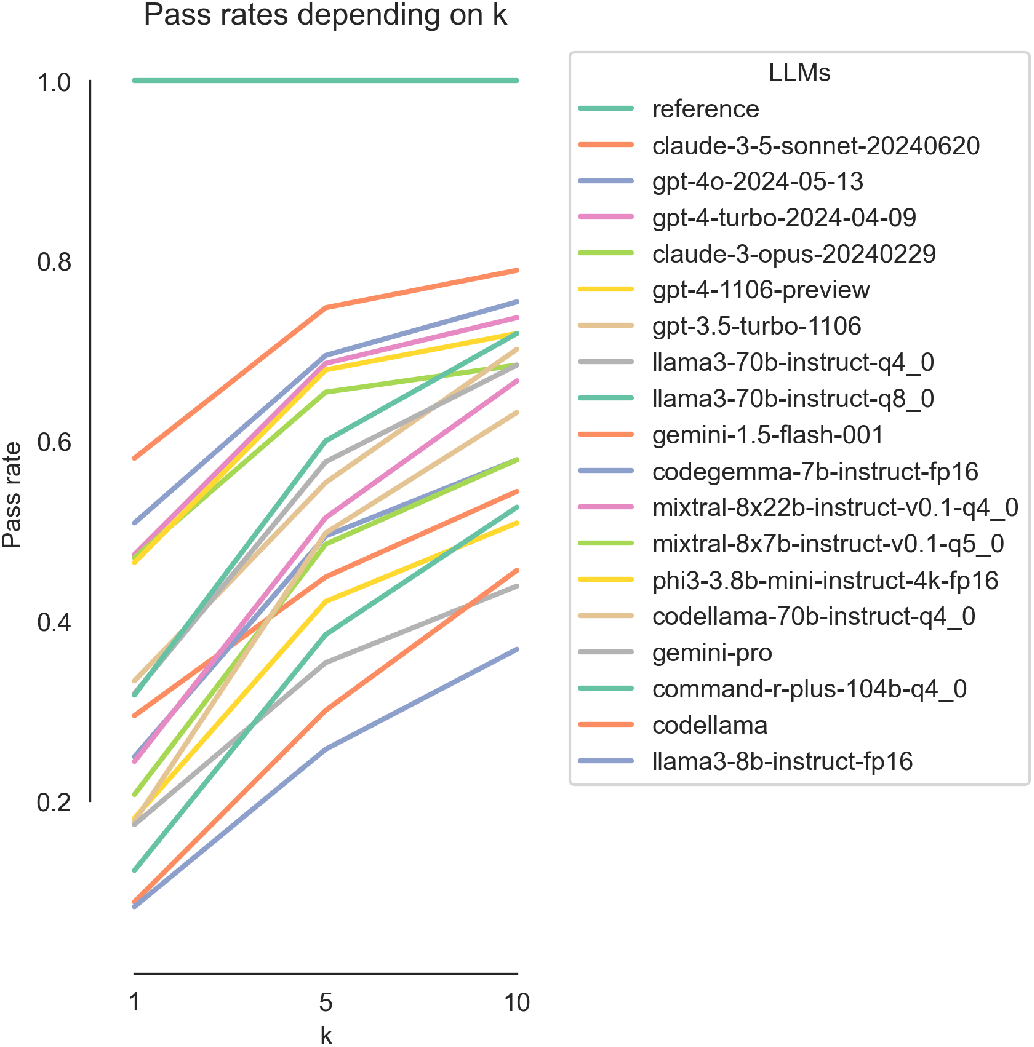
Detailed pass@k with *k* = 1, *k* = 5 and *k* = 10 is a way to estimate the chance to retrieve at least on functional code snippet when generating *k* samples. The corresponding, updated notebook is available online:https://github.com/haesleinhuepf/human-eval-bia/blob/main/demo/summarize_by_passk.ipynb

**Figure 4.**
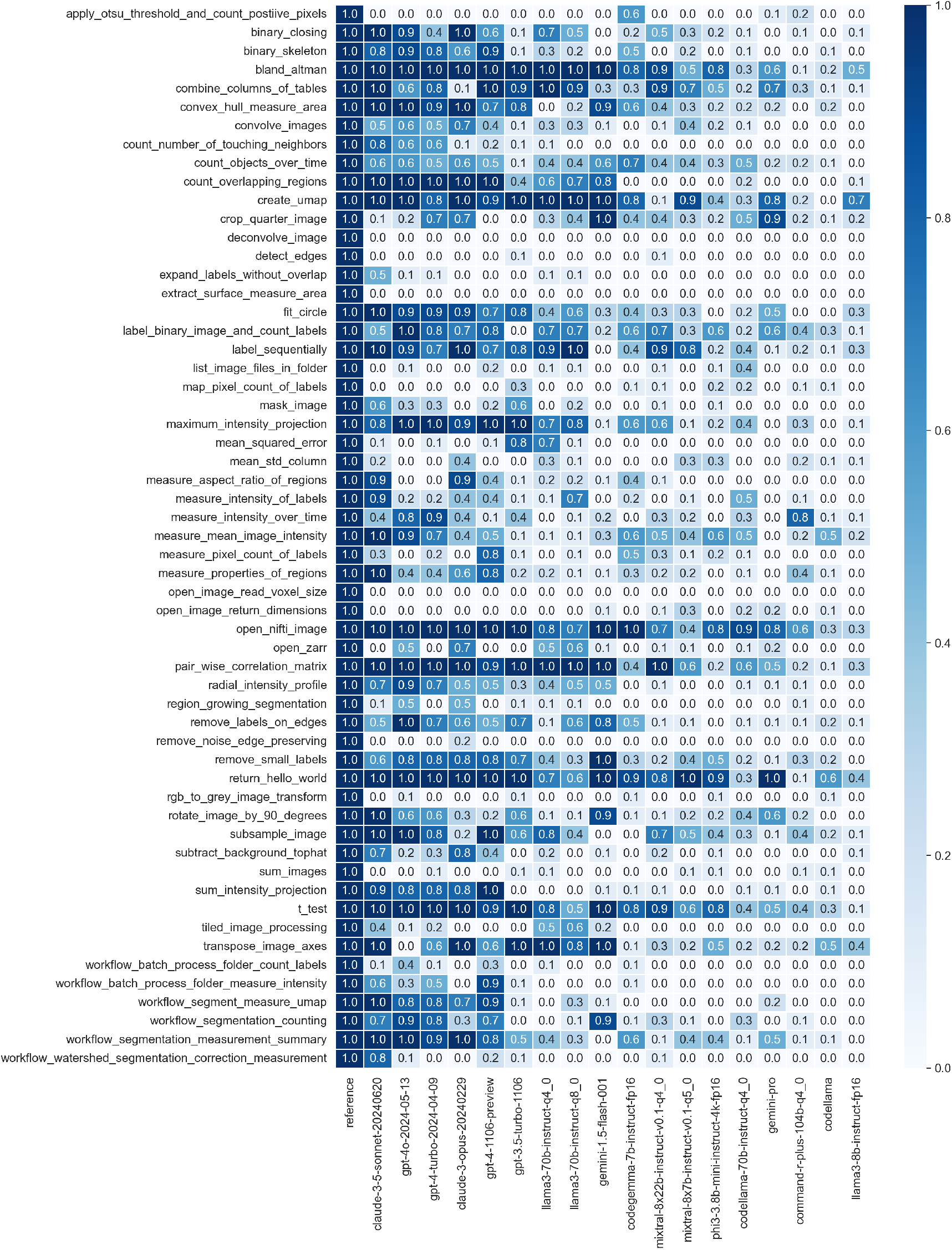
Test-cases and corresponding pass@1 for each LLM. Pass@1 reports the probability that a generated solution works if a user asks the LLM just a single time. The corresponding, updated notebook is available online:https://github.com/haesleinhuepf/human-eval-bia/blob/main/demo/summarize_by_case.ipynb

Details about how often the LLMs required specific Python libraries are summarized in Figure 5. For example, the skimage library was used in 22 of our human-written reference codes and thus, appears 220 times. By contrast, skimage was only used in a range of 68 to 154 generated code samples. Interestingly, our human-written reference codes were not using opencv, the “cv2” Python package, but the number of LLMs generated code where “cv2” appeared ranged from 31 to 192 cases.

**Figure 5.**
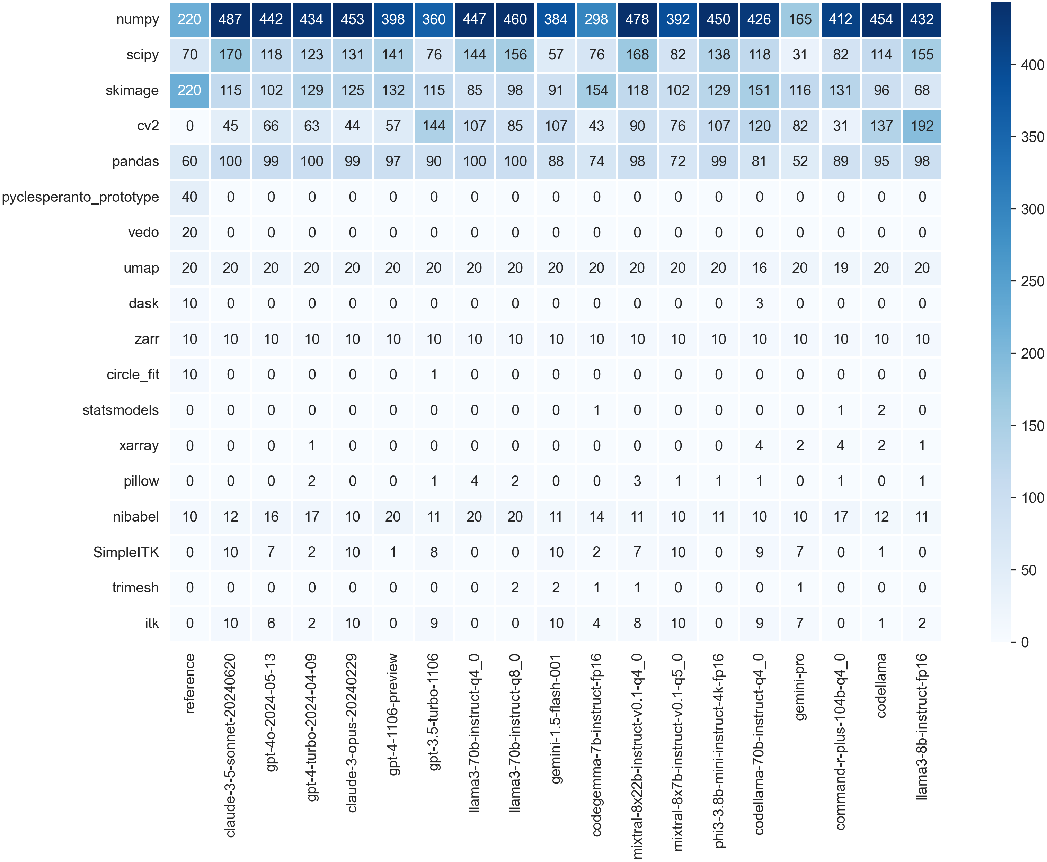
Used Python libraries in generated code from the tested LLMs. If one generated code snippet contained the same library twice, it is only counted once. The Notebook for generating this table can be found online:https://github.com/haesleinhuepf/human-eval-bia/blob/main/demo/summarize_used_libraries.ipynb

Common error messages and corresponding counts for each LLM are given in Figure 6. This analysis reveals a few systematic differences between the models, most notably gemini-pro often left out import statements leading to common error messages such as “name ‘np’ is not defined”. gemini-1.5-flash, a successor model did not show this pattern. llama3-8b and command-r-plus produced by far the most syntax errors.

**Figure 6.**
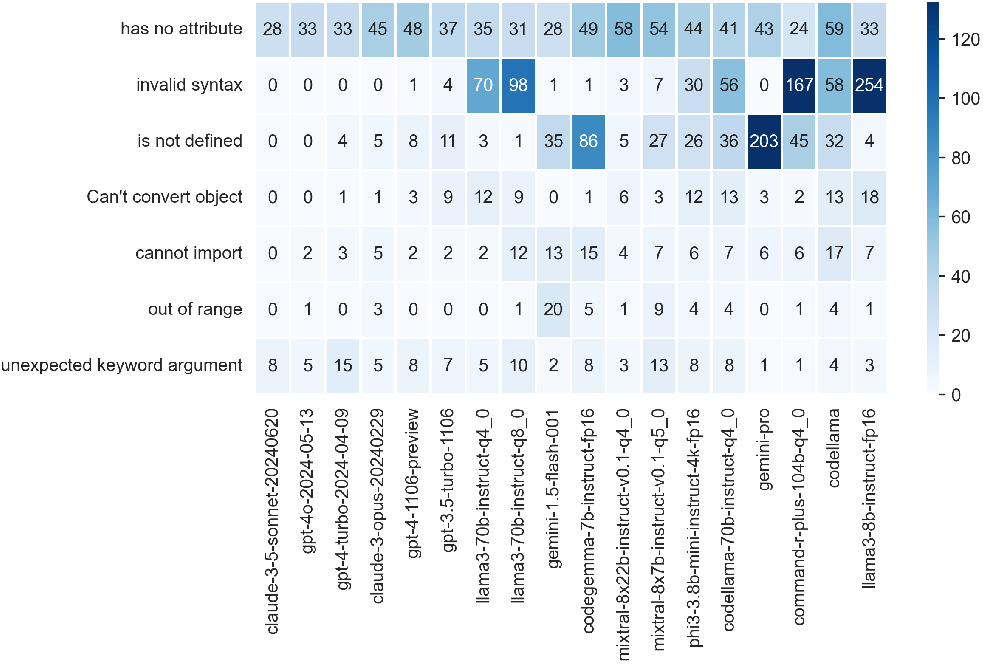
Snippets of common error messages and how often these were observed when evaluating the generated code from tested models. The Notebook for generating this table can be found online:https://github.com/haesleinhuepf/human-eval-bia/blob/main/demo/summarize_error_messages.ipynb

Sampling the LLMs using our prompts took from under 2 hours for the smaller models to 15-20 hours for the larger ones but exact comparisons aren’t available because different models ran on different hardware and infrastructure. The models gpt-3.5-turbo-1106, both gpt-4 models together, claude-3-opus-20240229, and claude-3-5-sonnet-20240620 caused costs of $0.52, $13.02, $24.58, and approx. $3, respectively. All other models did not cause direct costs as the use of their API was free.

## 4 Discussion

We presented a benchmark for comparing code generation capabilities of LLMs in the domain of Bio-Image Analysis. These benchmarks are crucial to decide, e.g. if and how to apply this technology in routine projects, training or advanced applications.

It should be mentioned that we did not use any code-completion tools, such as Github copilot, to write the test-cases. In general we think it is necessary to not use such LLM-based tools while writing the test-cases, because we might introduce a systematic bias towards the underlying LLMs. For example, Github copilot is based on the same model-family as ChatGPT. If the test-cases were written with the help of copilot, the benchmark might misleadingly reveal better performance of GPT-models.

Adding or modifying test-cases (e.g. the ones still failing), after a first benchmark has been executed, must be done carefully. We recommend a peer-review scheme, e.g. using Github pull-requests, to make sure good scientific practice is maintained. To this end, we provide a pull-request template with a short questionaire to support contributors. Additionally, we provide tools that detect if the code samples generated by LLMs are attempting to use Python libraries which are not installed yet. The missing libraries can then be added and the evaluation step can be repeated. We generally encourage contributors to add test-cases that can be implemented with common Python libraries, we however acknowledge that some subdomains (such as neuroimaging) may require specific libraries, e.g. nibabel [4], to interact with data. Mid- and long-term modifications should be done with care to maintain this benchmark. One of our major aims is to avoid biases towards specific LLMs.

We limited sample generation for the benchmarking to 10 samples per LLM per test-case. pass@k analysis was done using k=1, k=5 and k=10. The established standard set by [7] is 200 samples and k=1, k=10 and k=100. As our benchmark is in early development, we considered drawing 200 samples as not well-invested compute time and costs. Once the benchmark contains more test-cases and models, we will reconsider this decision. But also from a practical perspective when using LLMs on a daily basis, it appears unreasonable to draw hundreds of samples.

The estimated costs also demonstrate the potential of the technology. Requested code is commonly served within seconds, and drawing hundreds samples from the paid-per-prompt models causes relatively small costs depending on the model, so LLMs for BIA code generation can be cost-efficient. For situations were sending data to third parties isn’t an option, running LLMs locally is a viable option with decent performance from smaller models.

Our benchmark is a single-shot benchmark presenting the prompt to the LLM with no history of a former conversation. In daily use, one can interact with LLMs using chat-interfaces and iteratively engineer a prompt. Thus, it should be noted that the tested LLMs may be more capable than measured in our experiment, when used in a chat scenario. Furthermore, most of the models are generic LLMs but codegemma and codellama are specialized versions for coding. Interestingly, codegemma:7b performs as well or better as some much larger models suggesting that specialization can compensate for model size.

The test-case selection may introduce a certain bias from our subdisciplines. We often work with fluorescence microscopy imaging data, often showing nuclei, cytoplasm and membranes. Our test-cases are derived from practical situations we come across often. Mid-/long-term we hope that community contributions to the benchmark’s Github repository will allow us to cover the field more broadly. For example, algorithms analyzing histological, hyperspectral, or super-resolution imaging data, would be welcome additions to our test-case collection. On the other hand, we would exclude test-cases without practical relevance in bio-image analysis. For example, an algorithm for image-classification for natural images, e.g. showing cats and dogs, are considered off-topic and should not be included. We intentionally included test-cases and prompts which we presume are currently not solvable by LLMs, and we encourage the community to add more. With this, the benchmark could guide LLM developers in this field towards more advanced code-generation.

In our evaluation of LLMs we see two main groups of models: one group more capable than the other, as shown by pass-rates about a third as high as for the other models. There might be several reasons for this: 1) Many open-source models in our test are much smaller than the tested commercial models, e.g. llama3 has 70B parameters and the GPT models are about two orders of magnitude larger. Although model size limits LLM capabilities, it is interesting to note that specialist models like codegemma performs nearly on par with llama3 (and much better than codellama with a similar size) despite having an order of magnitude less parameters. 2) In bio-image analysis, we use some specific Python libraries, such as aicsimageio [5], vedo [18] or pyclesperanto_prototype [11], which might not be mentioned in the training data of some models. On the other hand, in natural image processing, libraries such as OpenCV [15] are common, while our community often uses scikit-image [26] for similar purposes. As natural image processing is a very active research field, the LLM’s training data may contain more examples from that domain. The DS-1000 benchmark [17], focusing on general data-science usecases, does not cover scikit-image or opencv. The focus of our benchmark may enable the LLM community to develop models covering more BIA use-cases and Python libraries, and thus help improve biological research.

We will continuously extend this benchmark with support of the BIA community. With the development of new models, we also would like to adapt the benchmark depending on how LLMs develop. For example, vision-models, LLMs that can also take images as input, need to be considered for benchmarking too. Our presented benchmark does not have the capability to test vision-models yet. Another direction for development could be efficiency of generated code as proposed earlier [9]. In particular in the context of processing 3D+t imaging data, accelerated image processing techniques can by key [10]. The presented benchmark could also be used to improve prompts systematically, for example in projects such as napari-chatGPT [23] and bia-bob [12]. More generally, it might be interesting to investigate different strategies for knowledge representation. For example, fine-tuned models, models with virtually unlimited context length, and retrieval augmented generation are three techniques for storing and accessing information how to use Python libraries. We would also like to evaluate these techniques quantitatively in the bio-image analysis context.

## 5 Conclusion

We developed a benchmark for measuring LLM performance in generating code for solving bio-image analysis tasks. It can guide experimentalists to decide which LLM to use and potentially to pay for when developing bio-image analysis scripts and tools. We also consider this benchmark for LLM-developers in our domain as a metric to guide further development. Last but not least: We encourage the community to send pull-requests with new test-cases to our Github repository to ensure the benchmark is covering needs in our field broadly. With this work, we want to establish a fully communitydriven approach to benchmarking LLMs for BIA.

## Acknowledgements

RH acknowledges the financial support by the Federal Ministry of Education and Research of Germany and by Sächsische Staatsministerium für Wissenschaft, Kultur und Tourismus in the programme Center of Excellence for AI-research “Center for Scalable Data Analytics and Artificial Intelligence Dresden/Leipzig”, project identification number: ScaDS.AI. NS acknowledges support by the BMBF (Federal Ministry of Education and Research) through ACONITE (01IS22065). JKH and CT thank EMBL IT services for their support with Kubernetes.

